# A comparative genome analysis of Rift Valley Fever virus isolates from foci of the disease outbreak in South Africa in 2008-2010

**DOI:** 10.1101/338251

**Authors:** Moabi R. Maluleke, Maanda Phosiwa, Antoinette van Schalkwyk, George Michuki, Baratang A. Lubisi, Phemelo S. Kegakilwe, Steve J. Kemp, Phelix A.O. Majiwa

## Abstract

Rift Valley fever (RVF) is a re-emerging zoonotic disease responsible for major losses in livestock production, with negative impact on the livelihoods of both commercial and resource-poor farmers in sub-Sahara African countries. The disease remains a threat in countries where its mosquito vectors thrives. Outbreaks of RVF usually follow weather conditions which favour increase in mosquito populations. Such outbreaks are usually cyclical, occurring every 10-15 years.

Recent outbreaks of the disease in South Africa have occurred unpredictably and with increased frequency. In 2008 outbreaks were reported in Mpumalanga, Limpopo and Gauteng provinces, followed by a 2009 outbreak in KwaZulu-Natal, Mpumalanga and Northern Cape provinces and in 2010 in the Eastern Cape, Northern Cape, Western Cape, North West, Free State and Mpumalanga provinces. By August 2010, 232 confirmed infections had been reported in humans, with 26 confirmed deaths.

To investigate the evolutionary dynamics of RVF viruses (RVFVs) circulating in South Africa, we undertook complete genome sequence analysis of isolates from animals at discrete foci of the 2008-2010 outbreaks. The genome sequences of these viruses were compared with viruses from earlier outbreaks in South Africa and in other countries. The data indicates that one 2009 and all the 2008 isolates from South Africa and Madagascar (M49/08) cluster in Lineage C or Kenya-1. The remaining of the 2009 and 2010 isolates cluster within Lineage H, except isolate M259_RSA_09, a probable segment M reassortant.

**Author summary:** A single RVF virus serotype exists, yet differences in virulence and pathogenicity of the virus have been observed. This necessitates the need for detailed genetic characterization of various isolates of the virus. The RVF virus isolates that caused the 2008-2010 disease outbreaks in South Africa were most probably reassortants. Reassortment results from exchange of portions of the genome, particularly those of segment M. Although clear association between RVFV genotype and phenotype has not been established, various amino acid substitutions have been implicated in the phenotype. Viruses with amino acid substitutions from glycine to glutamic acid at position 277 of segment M have been shown to be more virulent in mice in comparison to viruses with glycine at the same position. Phylogenetic analysis indicated the viruses responsible for the 2008-2010 RVF outbreaks in South Africa were not introduced from outside the country, but mutated in time and caused the outbreaks when environmental conditions became favourable.

## Introduction

Rift Valley fever (RVF), a mosquito-borne viral disease, affects humans and some species of ruminants including sheep, cattle, goats and buffalos. The causative agent, Rift Valley Fever virus (RVFV), belongs to the genus *Phlebovirus* in the family *Bunyaviridae*. The disease in livestock is characterized by abortion storms and high mortality of young animals. In humans, it manifests as febrile illness, resulting in retinal degeneration, severe encephalitis, haemorrhage; fatal hepatitis occurs in less than 1% of patients [1]. Protection of animals from the disease can be conferred by vaccination; however, there is currently no approved vaccine for use in humans. Transmission of RVF can be prevented by eliminating the mosquito vectors and avoiding human contact with infected tissues of infected animals.

The RVF virus has been isolated from more than 30 species of mosquitoes, belonging to at least six genera (*Aedes*, *Culex*, *Anopheles*, *Eretmapodites*, *Mansonia* and *Coquillettidia*). In a number of mosquito species, the virus has been isolated from both the insects and their eggs, suggesting different modes of transmission [2, 3]. Furthermore, the virus has been isolated from unfed mosquitoes reared from eggs obtained during inter-epidemic periods in Kenya and South Africa [3,4]. Eggs from floodwater mosquitoes can remain viable for several years, hatching during conducive climatic conditions associated with periods of high rainfall. Changing climate and farming systems can create conditions favorable for mosquito breeding, resulting in unexpected outbreaks of the disease [5].

The disease is endemic in eastern and southern Africa, but outbreaks have been reported in Egypt, Madagascar, Mauritania, Saudi Arabia, Sudan and Yemen [6,7,8]. No significant antigenic differences have thus far been demonstrated among isolates of this virus from different geographic locations, confirming the existence of a single RVFV serotype. Despite the single serotype, differences in virulence and pathogenicity of the virus have been observed, necessitating the need for detailed genetic characterization of the various isolates [9,10].

The genome of RVFV, like that of the other *Bunyaviridae*, consists of three-segmented, single-stranded negative- and ambi-sense RNAs with a total size of 12kb. The L (Large) segment codes for the viral RNA polymerase. The M (Medium) segment encodes a single precursor protein which is cleaved to produce the envelope glycoproteins G1 and G2, and two non-structural proteins of 78kDa and 14 kDa. In contrast, the ambisense S (Small) segment codes for the nonstructural protein NSs in the genomic sense and the nucleocapsid protein N in the antigenomic sense [9,11,12,13].

Because of its segmented structure, the genome of RVFV is thought to undergo recombination through reassortment, thereby contributing to its evolutionary dynamics [11,14]. In general, the RVFV genome is characterized by low genetic diversity (∼5%); consequently, it is difficult to statistically detect intragenic recombination events [9,15]. Similar to other arboviruses, all the genes of RVFV are under purifying selection and have evolved at distinct rates by accumulating mutations at 1.9 × 10^−4^ to 2.5 × 10^−4^ substitutions per site per year [9,14]. The previously estimated time to most recent common ancestor (TMRCA) is at around 124 to 133 years. This coincides with the importation of highly susceptible European breeds of cattle and sheep [10] into East Africa where the disease was first reported [15].

Despite the high sequence identity, nucleotide sequences of partial M segments from viruses isolated over 60 years from various countries have been grouped into 15 lineages [16]. The phylogenetic trees constructed from the nucleotide sequences of the three partial genome segments suggested the existence of reassortment, specifically of a 2010 isolate from a patient in South Africa. The individual was accidentally co-infected with live RVF animal vaccine and a RVF virus in lineage H [16]. Around this time, a RVF outbreak with unusual clinical presentation in animals was observed in South Africa. This outbreak had two distinguishing features: first, it occurred atypically in the absence of abnormally high rainfall; secondly, in addition to causing abortions storms, it had a high mortality among pregnant adult cattle [17].

The first case of RVF in South Africa occurred in the summer of 1950-1951 in animals and subsequently it was diagnosed in humans in 1951 [18, 19]. Three major outbreaks of the disease occurred in South Africa in 1950-1951, 1974-1976 and, most recently, in 2008-2011. There were minor incidents in the inter-episodic periods interspersing these outbreaks [20].

Confirmation of suspected cases of RVF in animals in South Africa is normally done at Agricultural Research Council–Onderstepoort Veterinary Research (ARC-OVR). Over time, the institute has accumulated a large collection of RVFV isolates from a majority of reported cases of the disease in South Africa. In order to obtain comprehensive information on the genetic composition of the RVF viruses (RVFVs) circulating in South Africa, we performed full genome sequence analysis of some of the viruses isolated from animals at discrete foci of the outbreaks which occurred during the 2008-2010 period. The genome sequences of these viruses were compared with the genome sequences of other RVFVs from earlier outbreaks in South Africa and other countries where the disease has occurred.

## Methods

### Growth, isolation and purification of RVF virus isolates

Presence of RVF virus nucleic acids in samples collected from animals suspected to be affected by the disease was confirmed by real-time PCR, using a slight modification of an established method [21].

Optimum conditions for efficient infection of Baby Hamster Kidney (BHK 21) cells (obtainedbfrom AATC) with RVF virus were established empirically using isolate M35/74, the challenge strain of RVFV [22]. The BHK 21 cells were grown in DMEM-F12 supplemented with 5%FBS (LONZA) and 1% pen/strep Amphotericin B (LONZA). These conditions were applied to infect BHK cells at a MOI resulting in the highest viral load. The infected cells were pelleted by centrifugation at 2 500 rpm for 5 minutes and the supernatant recovered. The supernatant was committed to sequence independent single primer amplification (SISPA) [23]. Briefly, the viral particles in the supernatant were treated with 100U DNase I and 4μg RNase at 37°C for 2h to remove possible host nucleic acids contamination. Viral RNA was extracted using TRIZOL LS kit (Invitrogen) according to the procedure provided by the supplier (Invitrogen). The RNA was recovered and used as the template in the first strand cDNA synthesis primed with FR26RV-N (5’GCC GGA GCT CTG CAG ATA TCN NNN NN3’ [23]. The single-stranded cDNA was the template for double-stranded cDNA synthesis using random 20mer primers and Klenow fragment of *E. coli* DNA polymerase. These products were subjected to PCR amplification using the 20-mer region of the above primer (FR26RV: 5’GCC GGA GCT CTG CAG ATA TC3’) in a reaction incubated in a thermocycler programmed to denature at 94 °C, 2 min then 35 cycles of 94 °C, 30 sec; 55 °C, 30 sec; 68 °C, 30s; with a final extension at 68 °C, for 10 min. The SISPA products were resolved by electrophoresis in 1% agarose gels.

### Construction of cDNA libraries

The SISPA products ranging in size from 0.2kb to 1.5kb were recovered from agarose gels and used in the preparation of library for sequencing reactions on the New Generation Sequencing (NGS) platforms exactly as described by the manufacturers (Roche Applied Science or Illumina). The sequencing was done on the Genome Sequencer 454 platform (GSFLX; 454 Life Sciences, Roche Applied Science; http://www.454.com); SISPA products from two random isolates were sequenced also on Illumina platform (http://www.illumina.com).

### Bioinformatics analyses of the sequence data

The sequence data obtained was processed and assembled into contigs using the appropriate software set to default values (Roche/454 Newbler for 454 Life Sciences Corporation, Software Release: 2.8 — 20120726_1306 or CLC Genomics Workbench, QIAGEN Bioinformatics).

The sequence data were subjected to further analyses using a combination of bioinformatics software. The nucleotide sequences were aligned using Clustal W [24] within the Molecular Evolutionary Genetics Analysis (MEGA) [25] set to optimum parameters for each sequence type. The best fitting nucleotide substitution model was determined for each genome segment using MEGA 6 and then applied in all the subsequent analyses. The aligned nucleotide sequences were used in calculating the mean pairwise distances and to derive phylogenetic trees using Maximum likelihood under 1000 bootstrap iterations [26].

Evidence for possible intragenic recombination events among the isolates was sought using different methods available from RDP3 [27]. Rates of molecular evolution for individual genome segments were estimated using Bayesian Markov Chain Monte Carlo implemented in the BEAUTI v1.8.1, BEAST v1.8.1, Tracer and FigTree packages [28]. The substitutions rates were estimated using both strict and relaxed uncorrelated lognormal molecular clock under General Time Reversible (GTR) model with gamma distribution (T4). The general Bayesian skyline coalescent prior was used and the MCMC allowed to run for sufficient number of generation with sampling every 1000 states, to ensure convergence of all parameters [28].

### Rift Valley fever virus genome sequence accession numbers

The nucleotide sequences of all the segments of the RVF isolates analyzed in the current study have been deposited in GenBank with accession numbers indicated in Table 1.

**Table I.**
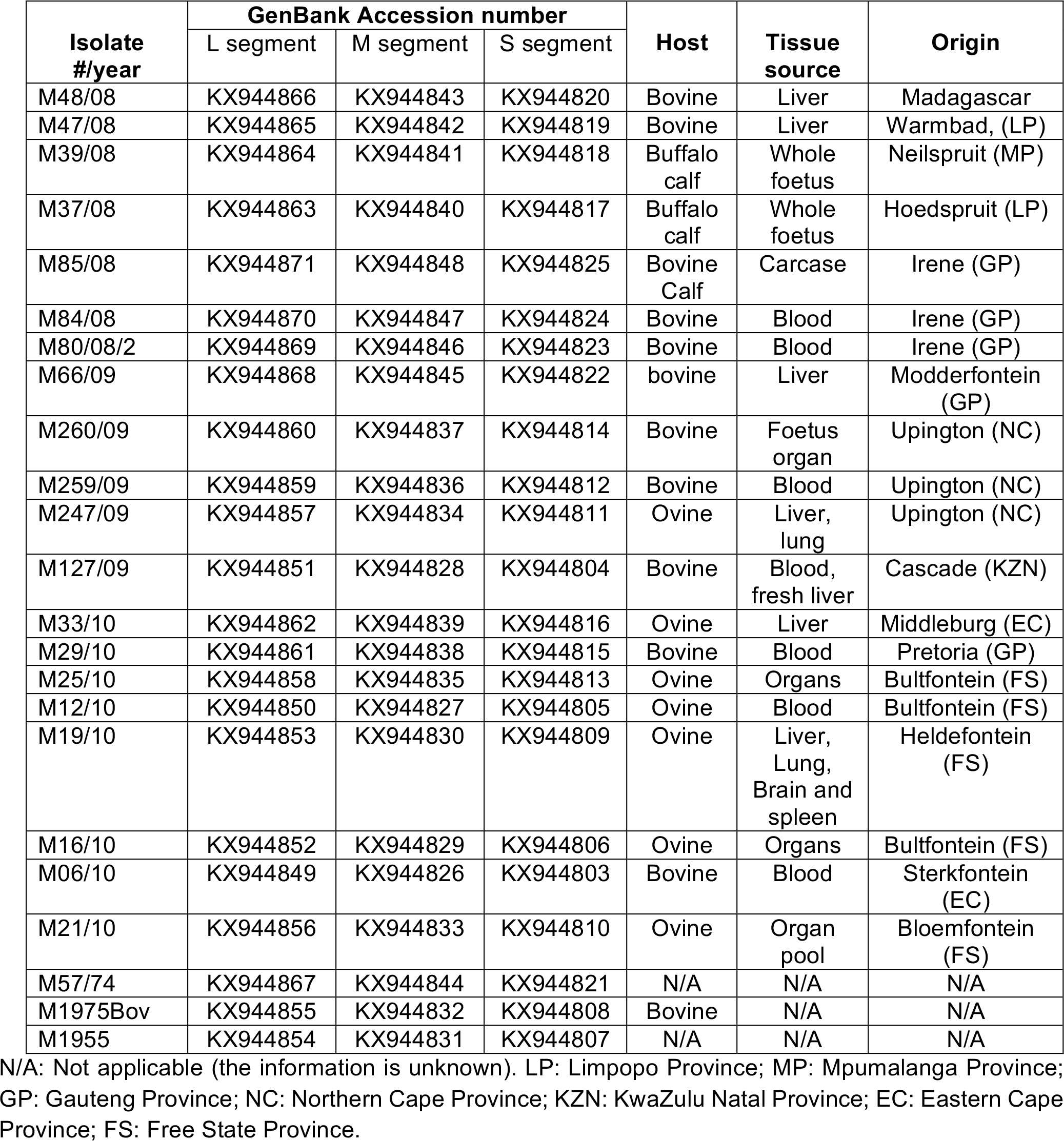
List of RVF virus isolates analyzed in the study and GenBank accession numbers assigned to the nucleotide sequences of their respective L, M and S segments. With the exception of Madagascar, origin imply the South African Province and nearest town from which the isolate originated.

## Results

Rift Valley fever virus SISPA products yield identical nucleotide sequence data irrespective of NGS platform used. This study focused on RVF viruses isolated during the disease outbreaks in South Africa in the period spanning 2008 to 2010, but also included viruses from the other major outbreaks in 1955 (M1955) and 1974-1975 (M57/74 and M1975Bov). The complete genome sequences of 23 isolates were determined using a combination of SISPA [23] and two independent NGS technologies. A representative profile of SISPA products obtained from the virus isolates is shown in Fig. 1. The 23 RVFV isolates whose genome sequences were determined represent four of the 15 reported outbreaks in 2008, three of the 19 outbreaks reported in 2009 and six of the 484 outbreaks in 2010 (Table 1) [29]. The complete genome sequence of isolate M48/08 from Madagascar was also determined (Table 1). The genomes of isolates M260/09 and M247/09 were sequenced on both GSFLX 454 and Illumina platforms, for comparison of sequence data obtained from SISPA products on either of these NGS technologies. The sequencing data obtained indicated that either of the technologies can be utilized with SISPA to obtain accurate genome sequences.

**Fig 1.**
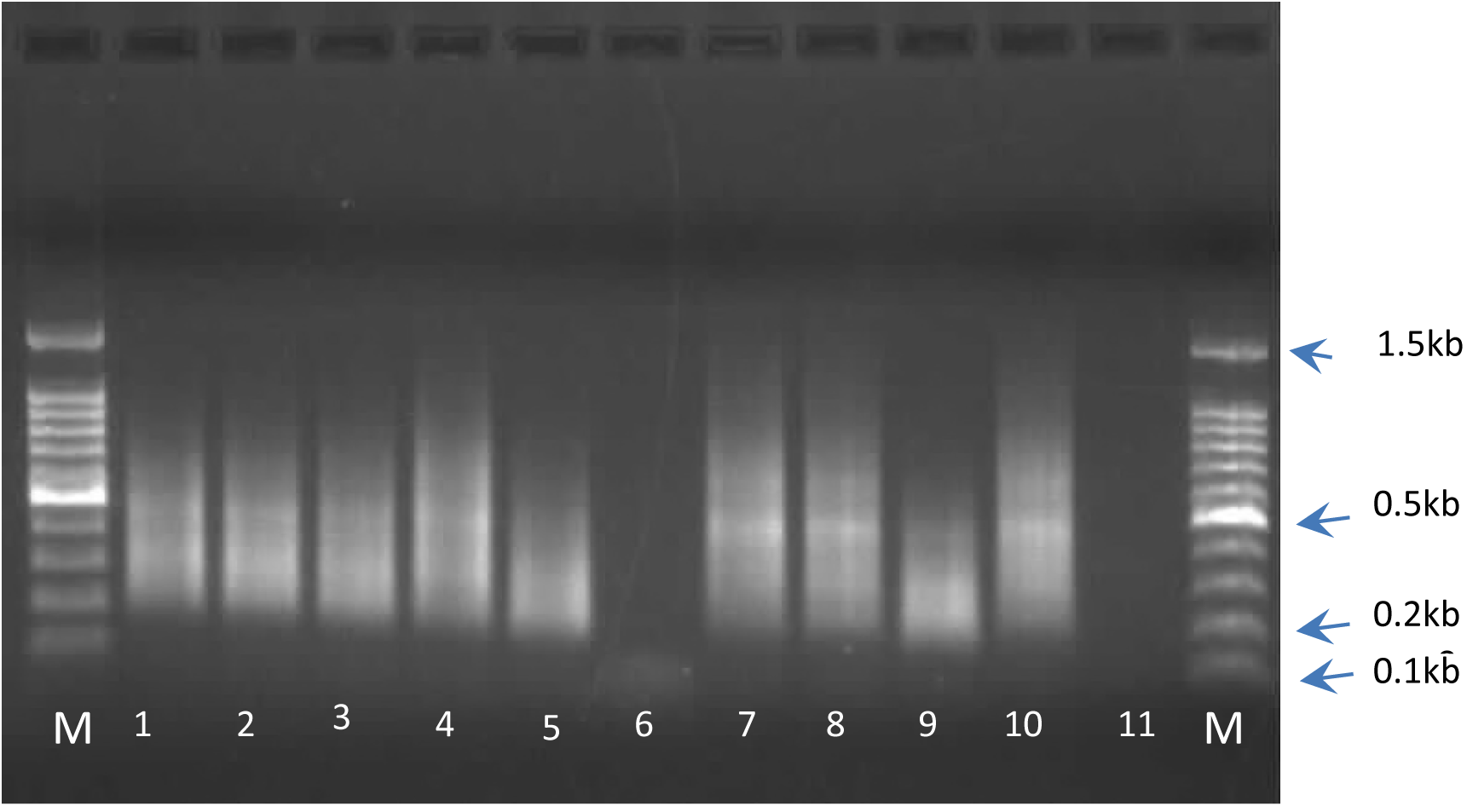
Photograph of a representative agarose gel in which SISPA products of RVF viruses were resolved. Lanes contain products as follows: lane 1: M03/10, lane 2: M15/10, lane 3: M06/10, lane 4: M19/10, lane 5: M21/10, lane 6: M22/10, lane 7: M23/10, lane 8: M25/10, lane 9: M26/10, lane 10: M33/10 lane 11: no DNA. Lanes labeled M contain DNA size markers, with corresponding sizes of some indicated in kilobasepairs (kb).

The nucleotide sequence data of all the RVFVs were assembled for complete S, M and L segments analyses and were deposited in GenBank with accession numbers as indicated in Table 1.

### Coding regions under selection pressure

Sequence alignments were generated for each of the three segments using all the available RVFV sequence data in GenBank. The alignments, which included full genome sequences from 120 – 140 isolates depending on the segment, were used in evaluating the evolutionary dynamics acting on each of the three segments.

Generally, sequence diversity among the segments were <5% among S or L segments, and <6% among M segments. Bayesian coalescent estimations of RVF genomes indicated that the segments evolve at a mean rate between 3.9 ×10^−4^ and 4.17 ×10^−4^ substitutions per site per year, regardless of the molecular model used. This is in agreement with previous Bayesian estimations [14,28]. Similarly, the estimated Time to Most Recent Common Ancestor (TMRCA) supports previous estimations of between 1880 and 1890 [14,28]. In order to determine the influence of substitution rate on biological function, we estimated the effect of differential selection pressures by calculating the rate of non-synonymous (d_N_) to synonymous (d_S_) substitutions. All the coding regions were found to be under purifying selection pressure (d_N_/d_S_ <1).

### Evidence for M segment reassortment but no intragenic recombination

Using Maximum likelihood trees, the phylogenetic relationship of the 23 RVFV isolates was assessed in relation to those of 50 other isolates genomes sequences of which were already in GenBank (S1 Table). Incongruences among the phylogenies of the individual genome segments were observed (Fig 2A-C), prompting the investigation into possible influence of recombination and reassortment.

**Fig 2.**
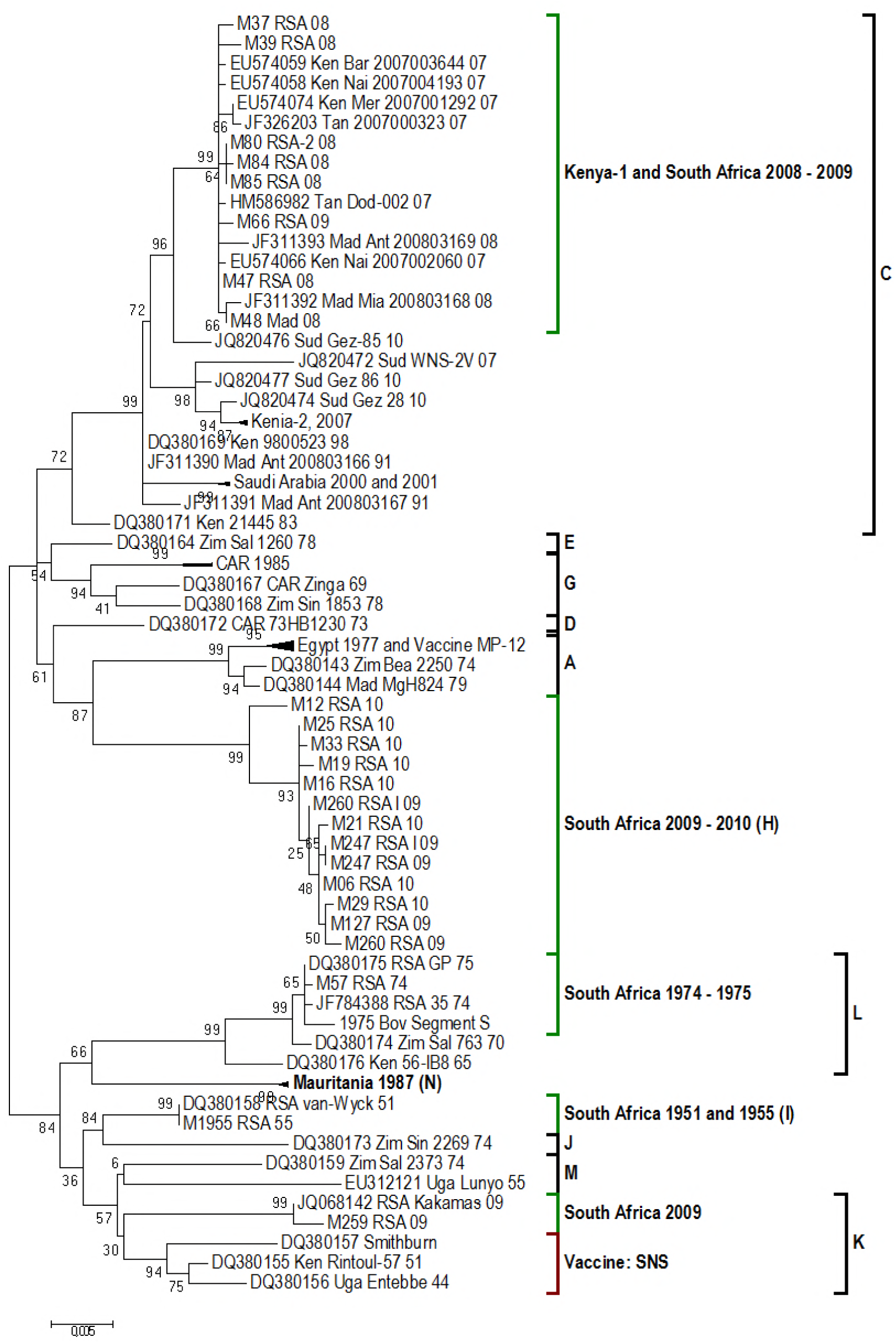
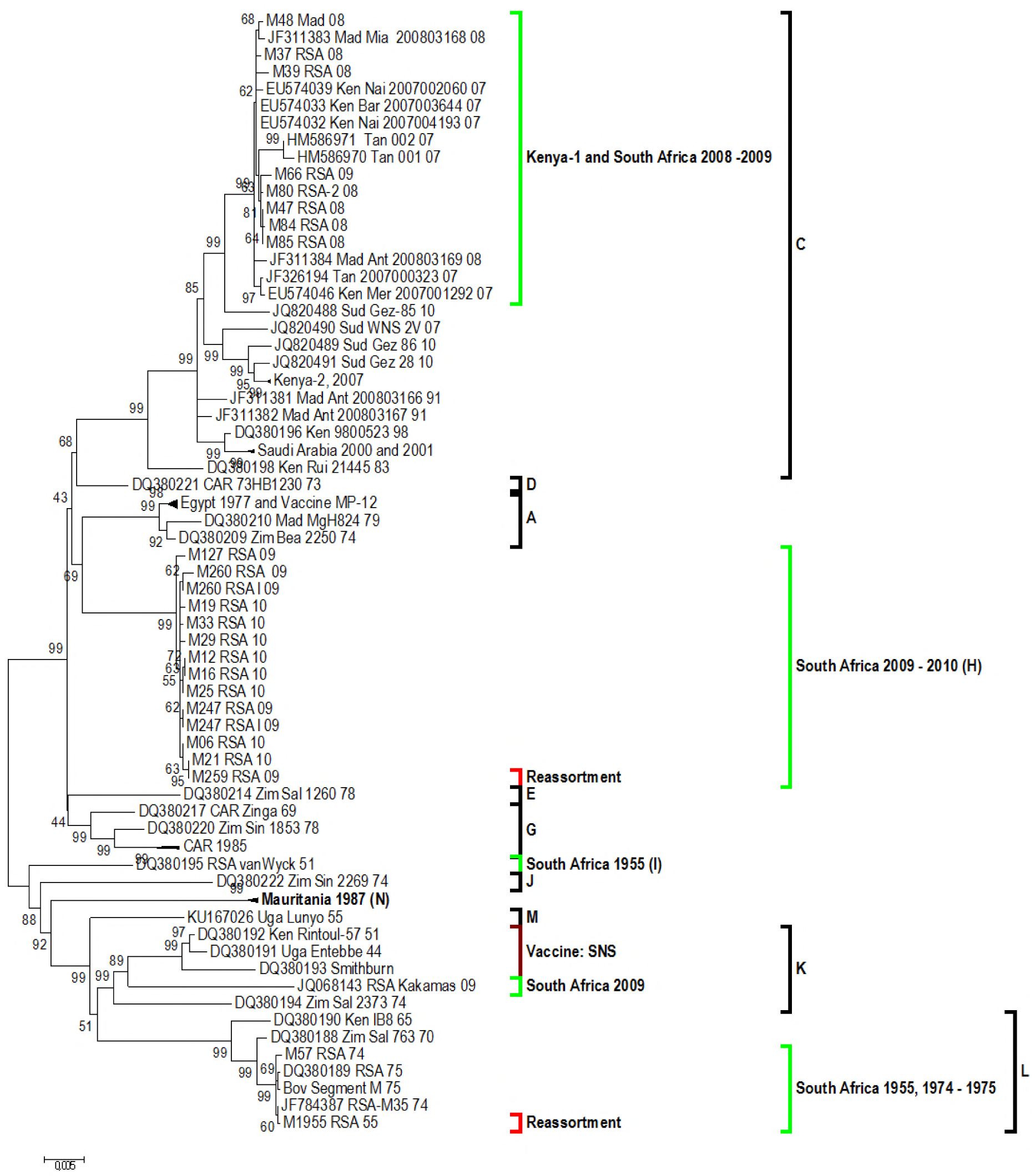
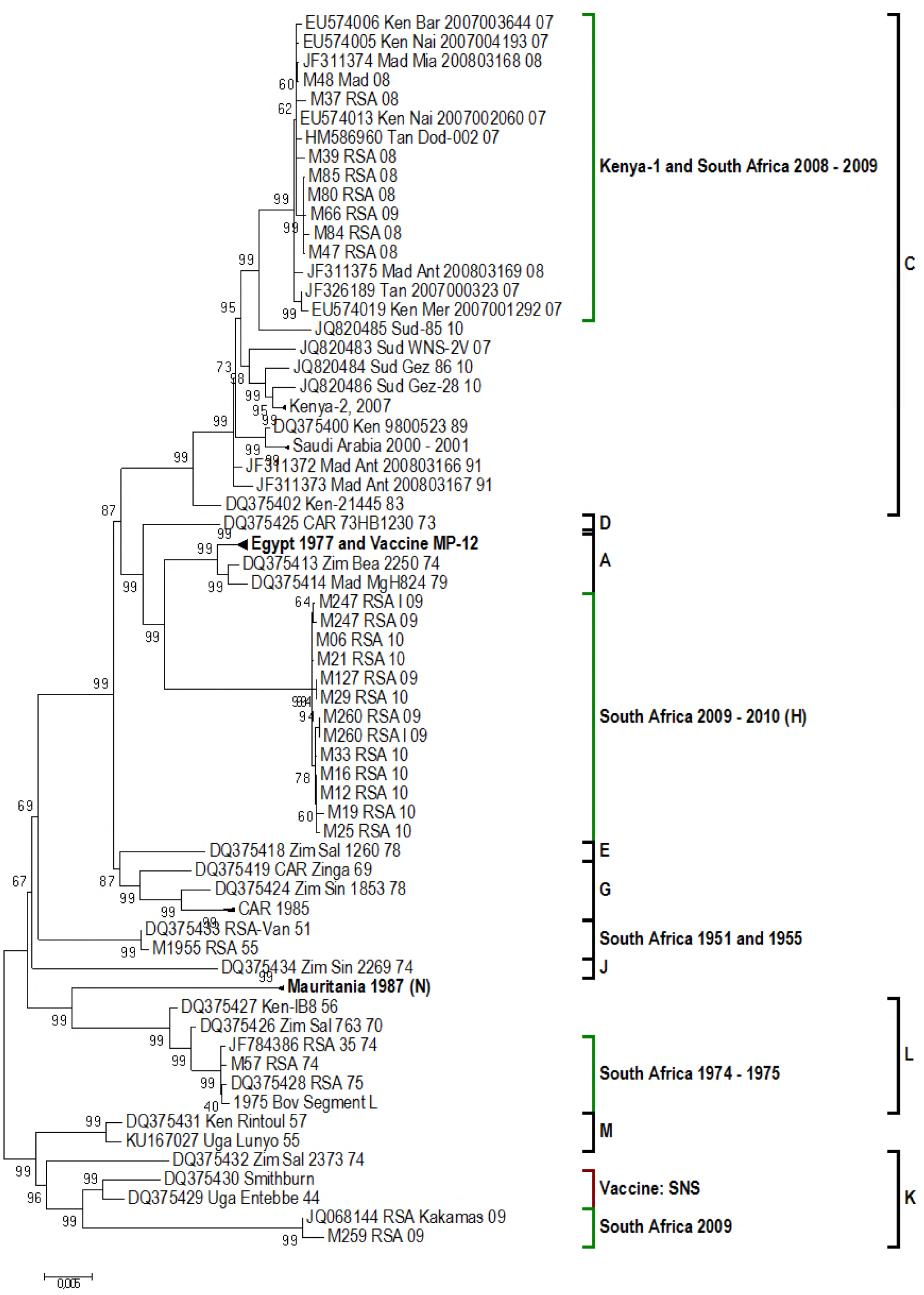
A-C. Phylogenetic trees derived from nucleotide sequence data of the three genomic segments of RVF virus segments. Segment S (A), Segment M (B) and Segment L (C) follows the same Lineage names as described by Grobbelaar et al., 2013. All Lineages containing RVF viruses from South Africa are presented in green. The two segment M re-assortments are indicated in red.

Insignificant statistical support for intragenic recombination events were predicted in all three segments (data not shown). This is explained by the low genetic diversity among the sequences [30]. Reassortment has been described for RVFV [11,14] and was therefore investigated utilizing the current data.

Fifteen lineages have been described using nucleotide sequences of parts of the three segments individually [16]. Our data indicate that one 2009 and all the 2008 isolates from South Africa and Madagascar (M49/08) clustered in Lineage C or Kenya-1[10,15] (Figs 2A, 2B and 2C). The remaining of the 2009 and 2010 isolates clustered within Lineage H [14], with the exception of isolate M259_RSA_09. The latter originated from serum of a bovine in Upington, Northern Cape Province, South Africa. Both its segments L and S cluster in Lineage K together with that of isolate JQ068143 from Kakamas (also in the Northern Cape Province); however, its segment M clustered in Lineage H along with the rest of the 2009 – 2010 isolates (Fig 2B). This indicates that isolate M259_RSA_09 is probably a segment reassortant from a coinfection with RVFVs in Lineages H and K. Whether this event occurred in an insect vector or an animal host Is not clear.

Segments L and S of isolate M1955_RSA_55 cluster in Lineage I together with the 1951 South African Van-Wyck isolate (DQ380158) (Figs 2A and C). However, segment M of this isolate (M1955_RSA_55) places it in Lineage L with the isolates of the 1974 and 1975 outbreaks in South Africa (Fig 2B). Thus, isolate M1955_RSA_55 was the second RVFV in this study, which had sequence features suggesting that it may be a segment reassortant.

Since both these putative reassortment events relate only to Segment M, which encodes two glycoproteins (Gc and Gn), the segment was subjected to additional analysis. The amino acid sequences of the glycoproteins encoded by the M segment of different RVF virus isolates from the 2008-2010 outbreak were compared to those previously published (S1 Table). The predicted amino acid residues are conserved with <3% sequence identity. Of the amino acid changes, 55% are conservative, 9.8% result in loss of a charge, 17% in gain of a charge and 2.7% in change of a charge. The positions of amino acid substitutions relative to the proportion of sequences with that change and those resulting in a change of charge are shown in Fig 3. Even though the majority of the substitutions are at the C-terminal region of the glycoprotein Gn, they are only observed in few of the sequences, the majority of them being conservative substitutions. One exception found was a change from D (Aspartic acid) to N (Asparagine) at the amino acid position 95, which is prominent in the 2008 – 2009 isolates in Lineage C, Fig 2B.

**Fig 3.**
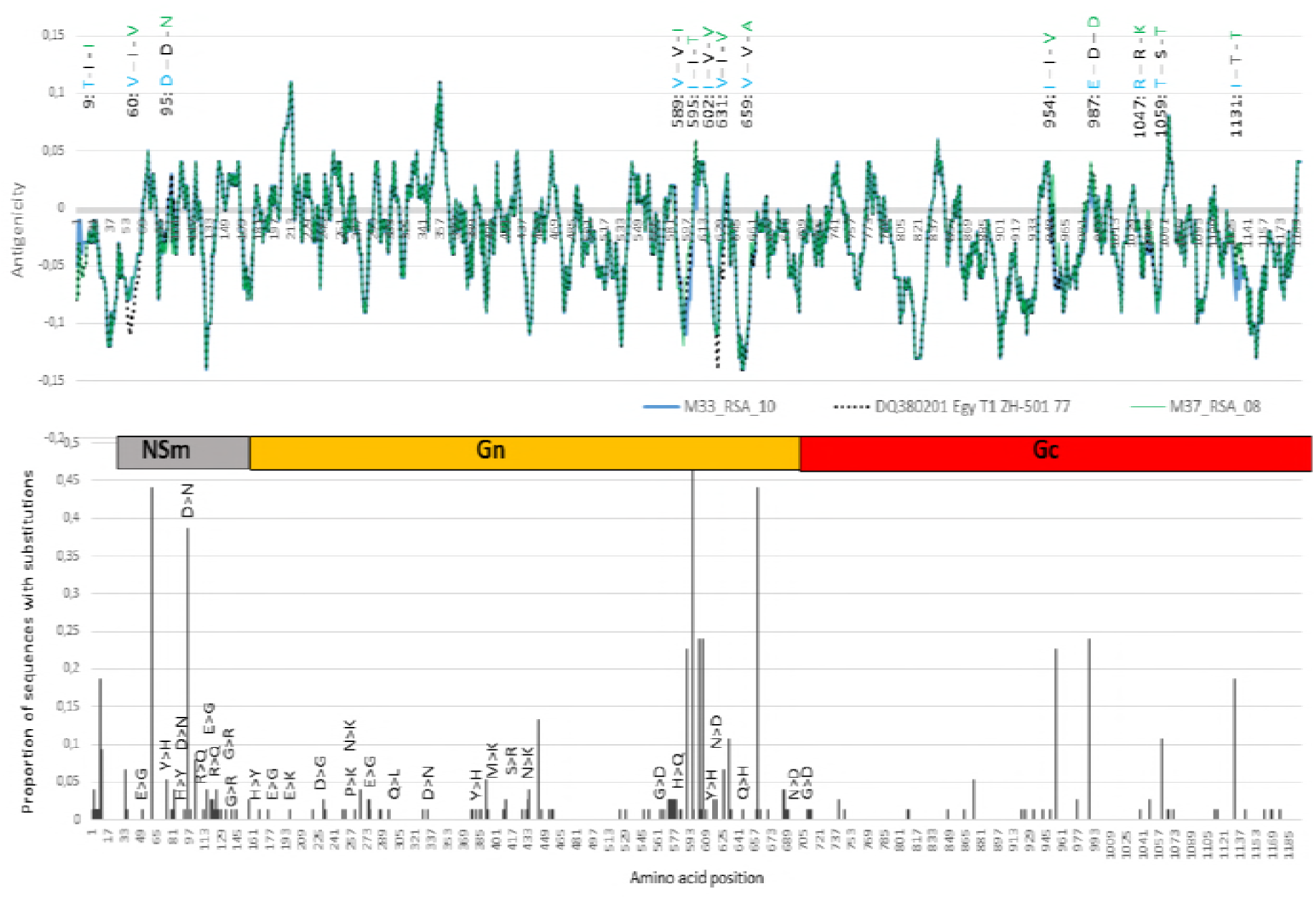
Welling antigenicity plots of proteins encoded by the M segments of isolates M33_RSA_10 in blue, ZH501-Egy-77 in black and M37_RSA_08 in green. Differences in amino acids residues among the three isolates are indicated on top of the antigenicity plots, with each isolate represented in its assigned colour. A graphical representation of the Non-structural protein (NSm), and glycoproteins (Gn) and (Gc) regions separate the antigenicity plots from the graph depicting the proportion of substitutions per amino acid position. These are representative of the 23 sequences of the M segments analyzed in this study and those of the previously published RVF viruses listed in S1 Table.

To investigate the possible influence of the amino acid changes on the antigenic properties of the viruses, we performed antigenicity predictions using Welling [32], with a window size of 11. Antigenicity plots for isolate M33_RSA_10, M37_RSA_08 and T1 ZH-501 isolated in Egypt in 1977 are shown in Fig 3. These isolates represent Lineages H, C and A, respectively. Virus ZH501 from the 1977 outbreak in Egypt has been shown to be associated with increased virulence in rats [33]. It was therefore included in this analysis for comparison with isolates from Lineages H and C [16]. The major differences in antigenicity predictions between ZH501-77 and the South African isolates are at positions 60 and 631 (Fig 3). Isolate ZH501-77 has a valine at both of these positions in contrast to the South African isolates which have isoleucine. Association of virulence with amino acid substitutions at positions 595, 631, 659 and 1059 has been shown in previous studies [10,33].

## Discussion

South Africa has experienced three major periods of RVF outbreaks, the first in 1950-1951, the largest in 1974-1976 and lastly in 2008-2011. In 2008 a total of 15 outbreaks were reported, localized to the central provinces of Limpopo, Mpumalanga, North West and Gauteng [29]. Complete genome sequence analysis of viruses isolated from the 2008 outbreak, clusters them in Lineage C together with isolates from a 2007 outbreak in Kenya, known as Lineage Kenya-1, Fig 2A, Fig 2B and Fig 2C [10,16]. In contrast to the centralized outbreak of 2008, 19 outbreaks which were reported in 2009 were distributed, with single cases in Mpumalanga and Gauteng, and the rest in KwaZulu-Natal, the Eastern Cape or along the Orange River in the Northern Cape [29,34]. Similar to the isolates from the 2008 outbreaks, the 2009 isolate from Gauteng clustered in Lineage C, within Lineage Kenya-1 (Fig 2A and B). This 2009 Gauteng outbreak appears to have been caused by the 2008 RVF viruses present in that region. Isolates from both the geographically distinct KwaZulu-Natal and Northern Cape outbreaks of 2009, clustered in Lineage H (Fig 2A). This lineage includes isolate M259_RSA_09, a segment M reassortant, whose segments L and S cluster with the 2009 Kakamas isolate (JQ068142) in Lineage K (Fig 2B). It is therefore possible that the RVF viruses associated with the majority of the outbreaks in 2009 originated from a single source.

In 2010 a total of 484 outbreaks were reported in every province of South Africa, except KwaZulu-Natal [20]. The initial report of an outbreak was from Bultfontein and Brandfort, both in the Free State and subsequently cases were reported from across the country [29]. Similar to the virus isolates from the 2009 outbreaks in KwaZulu-Natal and the Northern Cape, all the isolates from the 2010 outbreaks clustered in Lineage H (Fig 2A and B). The eight isolates from the 2010 outbreak analyzed in this study (Table 1 and Fig 2) are not necessarily statistically representatives of the 14342 cases reported in that year, but analyses of their nucleotide sequence data support speculations that the 2010 outbreak was a continuation of the 2009 in KZN and Northern Cape outbreaks. The clustering of isolates in lineage H (Fig 2A) gives an indication that new strains could evolve due to nucleotide substitution (Fig 3), albeit at slow/low rate. It is possible that these viruses were not introduced from elsewhere outside South Africa, but rather that they mutated over time and caused outbreaks when suitable conditions prevailed.

This study has contributed full genome sequence of RVFVs M57_RSA_74 isolated during the 1974 outbreaks and M1955_RSA_55 isolated from one of the 28 outbreaks in 1955 [29]. The largest RVF epidemic reported in South Africa were between 1973 and 1976, with mortality rates of 95% and cases reported from every province [29, 35]. Previous studies have clustered the 1973-1975 isolates into Lineage L along with a 1970 isolate from Zimbabwe and a 1956 isolate from Kenya[16]; as expected isolate M57_RSA_74 clustered with these (Fig 2A, 2B and C). In contrast, Segment S and Segment L of isolate M1955_RSA_55 clustered with a 1951 South African isolate known as van-Wyck in Lineage I (Fig 2A and C) [16], but Segment M clustered with isolates from the 1973-1976 outbreaks in Lineage L (Fig 2B), making this 1955 isolate a segment M reassortant. The occurrence of segment M reassortment in M1955_RSA_55 indicates that multiple RVF virus lineages can co-circulate, resulting in reassortant viruses re-emerging decades later causing disease outbreaks.

The evolutionary dynamics of RVFVs are characterized by low substitutions rates (3.9 ×10^−4^ and 4.17 ×10^−4^ substitutions per site per year) under strong purifying or negative selection with the major genomic diversity resulting from reassortment [11,14]. Similar evolutionary dynamics have been described in other arboviruses such as bluetongue virus and Epizootic haemorrhagic disease virus, due to the obligatory replication of the virus in both its insect vector and mammalian host [36]. The majority of reasortment events described in RVFV involves the exchange of segment M, resulting in antigenic shift due to the two glycoproteins Gn and Gc encoded by this segment [10,14,16].

Although RVF virus is antigenically homogenous, various isolates of the virus exhibit differences in virulence, evident upon infection of the mammalian host [33,36]. Whereas some of these differences may be attributable to the individual host, others are inherent to the virus. Differences in virulence and lethality of RVF virus isolates have been observed during the experimental infection of BHK cells [37], mice [33], sheep [38] and cattle [37]. Significant differences associated with the severity of RVF in humans have been observed [39, 40]. An increase in the severity of RVF since the 1977 outbreak in Egypt to the devastating outbreak during the 2006-2008 in East Africa have been observed [41]. A clear association between RVFV genotype and lethal phenotype has not been established; however, various amino acid substitutions have been implicated in this phenotype [37]. Therefore, indicative amino acid changes in some of the RVF proteins were investigated. The most prominent substitution in the glycoproteins are 595 I>V, 605 R>K, 631 I>V, 659 V>A located in Gn and 1059 S>T within Gc [10]. Another variation was identified in ZH501, isolate from a human in Egypt during an outbreak in 1977, which resulted in the change of Glycine to Glutamic acid at position 277. The virus with the Glutamic acid displayed an increased virulence in mice, compared to the virus with Glycine in the same position [33].

The majority of RVF viruses analysed in this study had Glutamine at position 277, except wild type isolate 763/70 from a foetus aborted during an outbreak of the disease in Zimbabwe in 1970 [10]. This study identified additional substitutions between the lethal isolate ZH501-77 from Egypt and isolates belonging to Lineage H from 2010 in South Africa (Fig 3B). The substitutions included 602 V>I, 987 D>E and 1131 T>I. The impacts of each of these substitutions on the pathogenicity of RVFVs remain to be investigated.

The fact that the sequence data of two isolates (M260/09 and M247/09) generated by different NGS platforms clustered together (Fig 2A) demonstrates that the two platforms produce identical sequences. One caveat with the dataset analyzed in this study is that the isolates might not be representative of the RVF viruses circulating during the 2008-2010 outbreak. This is inherent in the way the study was done: samples brought for testing at the ARC-OVR are opportunistic and are not necessarily representative of cases of RVF in animals in South Africa. During this period; the RVFVs whose genomes could be analyzed are the viruses that infect BHK 21 cells growing in culture media; and finally, good quality sequence data could not be obtained from all RVFVs, which were isolated in cell culture. A different picture of viruses and their potential quasispecies might emerge when the analyses are performed on viruses obtained directly from representative proven clinical cases. This is currently unattainable in our system, but determining the entire RVF viral genome sequence directly from clinical samples is being investigated.

## Acknowledgements

We thank Dr Marco Romito for performing PCR tests which identified the samples positive for RVF virus, the technical staff in Virology Laboratory for isolating RVF viruses from the positive samples; and Drs Keoagile Oliver Bezuidt and Oleg Reva, FABI, University of Pretoria, for assistance with some parts of the bioinformatics work.

Preliminary parts of this work have previously been presented at: Interregional Conference on Rift Valley Fever in the Middle East and Horn of Africa, 21 - 23 April 2015, Djibouti City, Djibouti and 11th Annual Sequencing, Finishing and Analysis in the Future (SFAF) conference, 1^st^ - 3rd June 2016, Santa Fe, NM, USA.

---

**Table S1.**
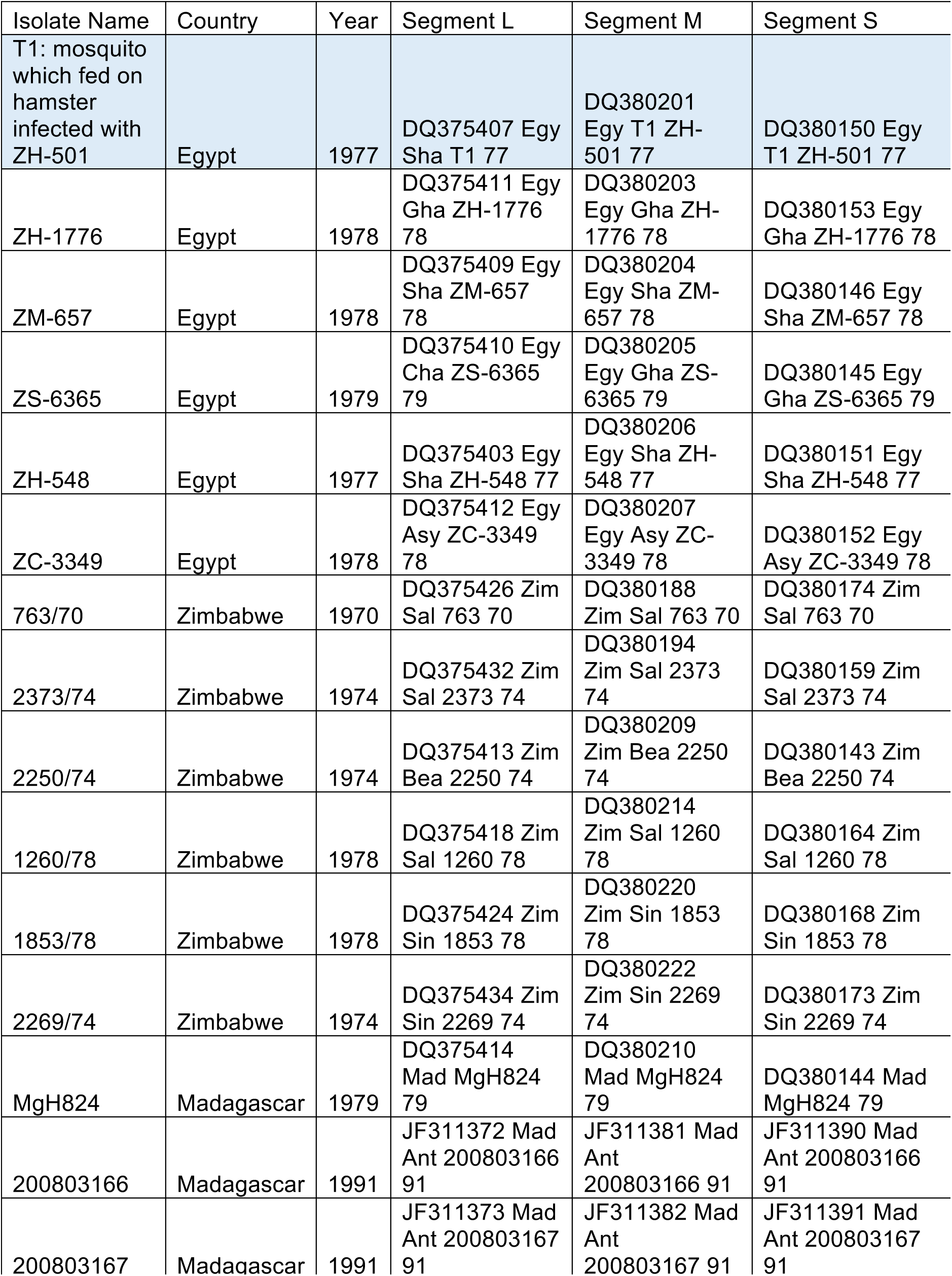
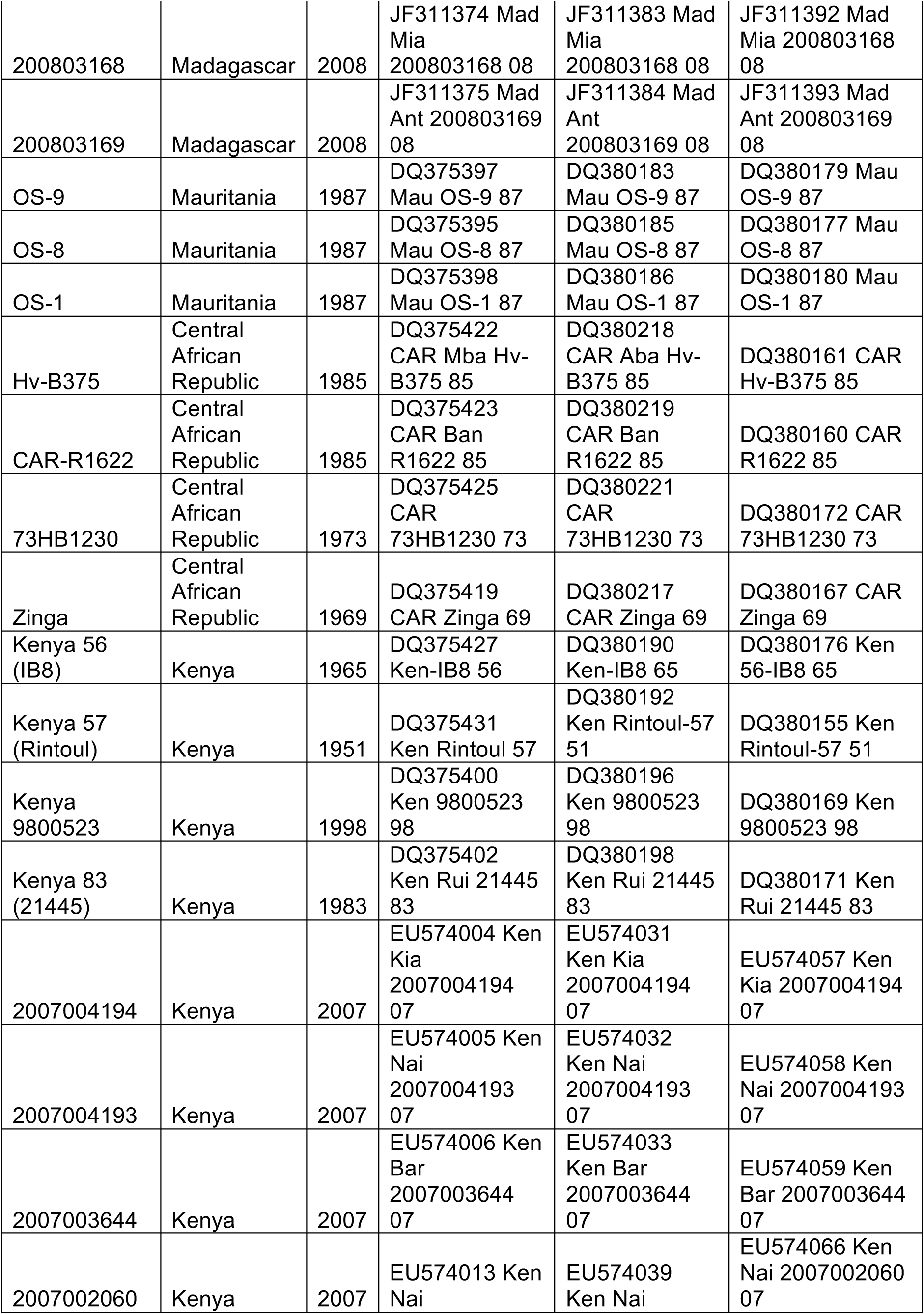
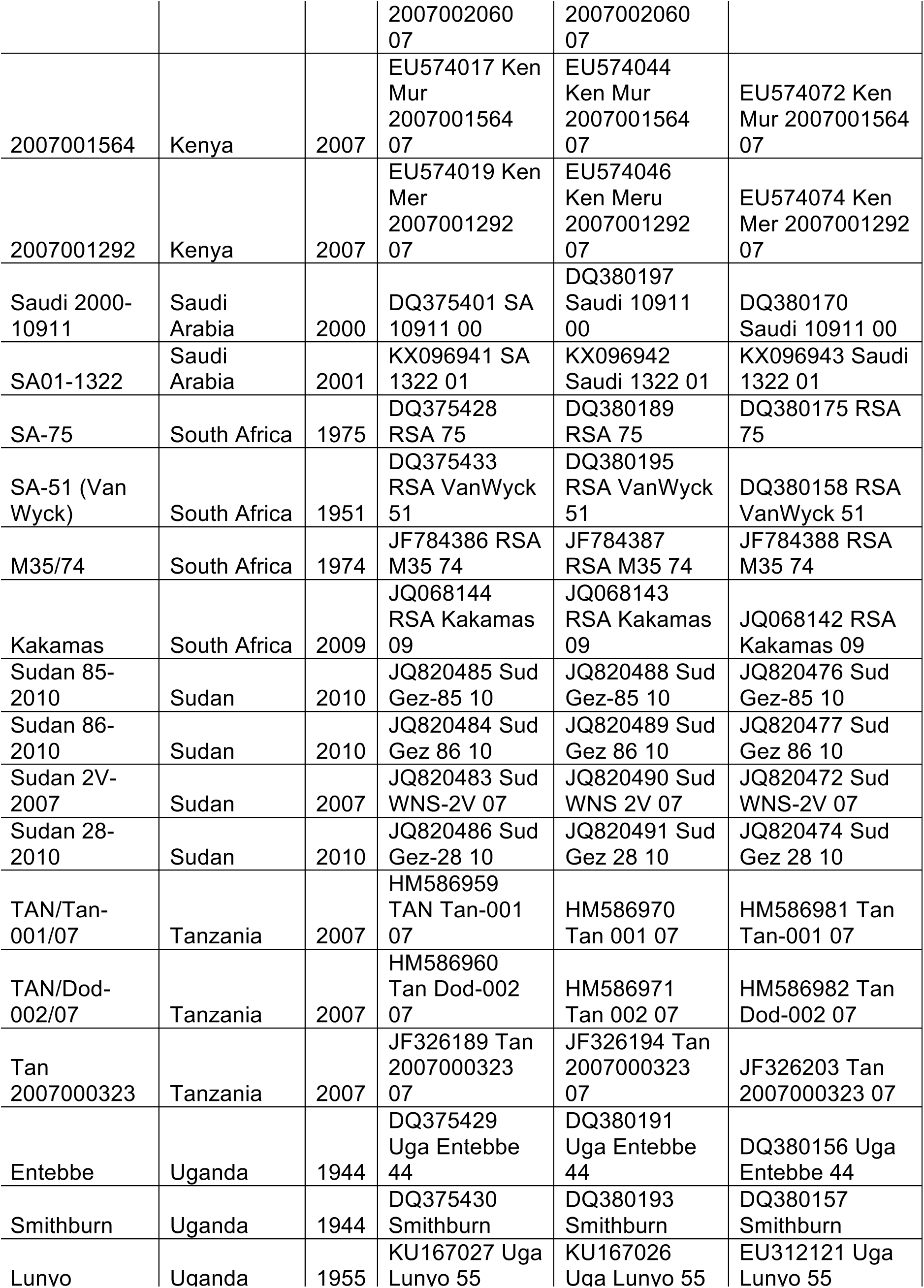
Previously published RVF viruses used in the study

